# Panache: a Web Browser-Based Viewer for Linearized Pangenomes

**DOI:** 10.1101/2021.04.27.441597

**Authors:** Éloi Durant, François Sabot, Matthieu Conte, Mathieu Rouard

## Abstract

**Motivation:** Pangenomics evolved since its first applications on bacteria, extending from the study of genes for a given population to the study of all of its sequences available. While multiple methods are being developed to construct pangenomes in eukaryotic species there is still a gap for efficient and user-friendly visualization tools. Emerging graph representations comes with their own challenges, and linearity remains a suitable option for user-friendliness.

**Results:** We introduce Panache, a tool for the visualization and exploration of linear representations of gene-based and sequence-based pangenomes. It uses a layout similar to genome browsers to display presence absence variations and additional tracks along a linear axis with a pangenomics perspective.

**Availability:** Panache is available at github.com/SouthGreenPlatform/panache under the MIT License.

## Introduction

The widespread use of fast and affordable sequencing technologies unveiled how much genomic information was lost when relying on a single and unique reference genome. For instance, it was found that about 10% of additional DNA was not captured by the current human reference genome (Sherman *et al.*, 2019). By leveraging data of multiple references instead, a new era of genomics emerged: Pangenomics. This is now being applied from bacteria to Eukaryotes and has been increasingly used in more complex genomes such as human and plants. As reviewed in (Golicz *et al.*, 2019), some studies handle this approach through a gene or functional annotation lens while others extend it to DNA sequences, especially when the studied organisms are eukaryotes.

Still, more tools are needed to help pangenomics to reach a broader audience within the scientific community (Computational Pan-Genomics Consortium, 2018; Golicz, Batley, *et al.*, 2016; Tranchant-Dubreuil *et al.*, 2019). While recent progress has been made to compute and store pangenomes (Garrison *et al.*, 2018; Li *et al.*, 2020), the tool landscape is particularly barren when it comes to visualization. Tools such as Pan-Tetris (Hennig *et al.*, 2015), PanViz (Pedersen *et al.*, 2017) or PanX (Ding *et al.*, 2018) were designed for genebased pangenomes and do not scale well to large-scale eukaryotic studies. Indeed, this gene-centric definition does not take into account positions within genomes, thereby blending paralogs together, and ignoring non-coding sequences despite their crucial influence on phenotypes (Maston *et al.*, 2006). The current trend for sequence-based pangenomes is to use graph visualization software like Bandage (Wick *et al.*, 2015), a general tool for navigating assembly graphs, but alternatives dedicated to pangenomes and their inner properties are yet to be refined and adopted. For instance Sequence Tube Maps (Beyer *et al.*, 2019) and MoMi-G (Yokoyama *et al.*, 2019) both focus on structural variations from individual genomes but lack information on the pangenome itself. As an example, they do not have direct visual cues for the identification of the most represented parts of a pangenome.

As useful as graph representations may be, they can easily be overloaded with content, resulting in a ‘hairball’ effect that is hard to read and explore (Yoghourdjian *et al.*, 2020). Linear representations of genomes have their own weaknesses (Nielsen and Wong, 2012) but are widely used in a variety of genome browsers, and most efficient when it comes to exploration tasks. Here we introduce Panache, the PANgenome Analyzer with CHromosomal Exploration, a web browser-based viewer which renders interactive linear representations of pangenomes one panchromosome at a time.

## Features

Panache is designed to display a linear representation of pangenomes. This representation is based on the idea that every genome within a pangenome can be divided into multiple blocks, with each block potentially shared with other genomes. Such blocks could be either DNA sequences (extracted from the nodes of a graph pangenome for example) or genes. Blocks from all genomes could be laid out and ordered along a single string, which would then serve as a flattened pan-reference, as illustrated in Figure 1A. One could also imagine ordering the blocks according to an existing genome instead, using its own linear coordinate system as a reference. Every pangenomic block can therefore be represented in an easy-to-browse visualization, with additional tracks of summarized information such as a block presence/absence status or whether it belongs to the core genome (most present blocks) or the variable genome (also referred to as dispensable genome).

**Figure 1.**
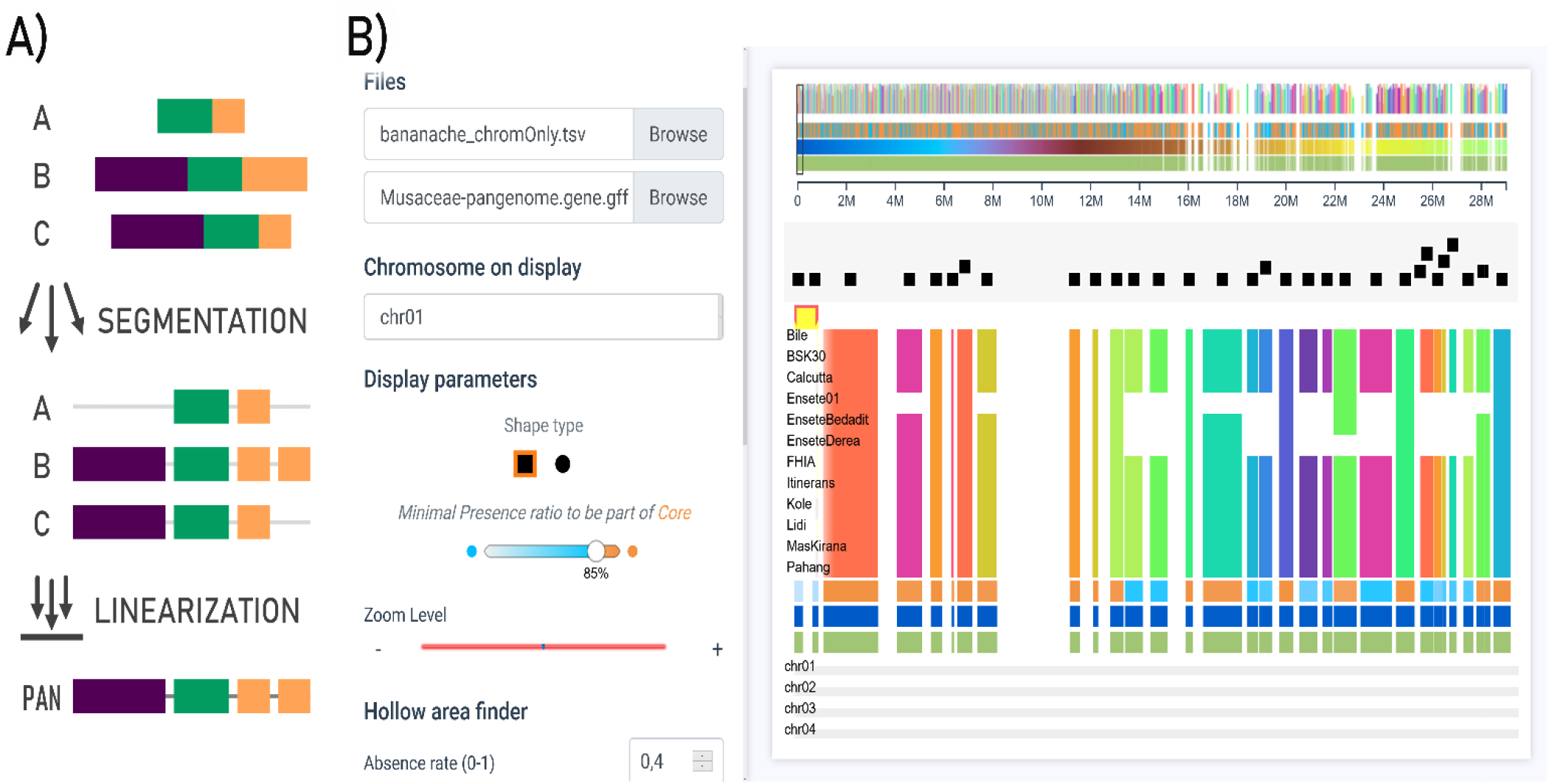
Panache offers a linear representation of pangenomes, with blocks information detaile through multiple tracks much like classic genome browsers. **A)** Linearized pangenomes represent chains of present/absent pangenomic blocks on a single string. **B)** Panache’s interface for browsing through the Presence/Absence matrix and navigating through panchromosomes.

While a graph representation might give a better sense of structural variations or of a genome’s full sequence within a pangenome, a linear representation allows more reproducibility when exploring data thanks to its fixed and ordered coordinate system. Moreover, missing information can be visually hinted even when not directly available. For example, an additional track can specify which blocks are repeated elsewhere in the pangenome.

The tool comes with a variety of navigation related functionalities: choosing which panchromosome to display and browsing through it, modifying display size, jumping automatically to areas enriched in absent blocks. It also allows modifications to the core/variable threshold and hovering over visual elements to display additional information such as functional annotation pop-up windows. All available functionalities are further detailed within Panache’s documentation.

Figure 1B illustrates how Panache displays linear pangenome data using a pangenome generated in banana (Rijzaani *et al.*, 2021). A set of 34,878 genes from 15 genotypes have been grouped into chromosomes and positioned linearly on a panreference. Here a user can quickly identify lines with missing genes, and which genes belong to the core genome (in orange) or to the variable genome (in blue). Details about the individual gene annotations can be accessed by hovering over the beeswarm plot on top of the presence/absence matrix.

## Implementation

Panache is a JavaScript application built with Vue.js 2 and additional librairies, namely D3.js v5 (enabling linkages between data and SVGs) and Vuex. For easy deployment, we have created a docker instance that can run Panache through nginx but the production files are available for deployment through other means.

Panache can take two types of files as input. The main file is a presence/absence matrix file in a BED-like format. Each line stores information about one pangenomic block (either a gene or a sequence) detailed through multiple columns, starting with linear position data and ending with the presence/absence information within every genome. An optional GFF file of annotations on the linear pangenomic coordinates may be loaded in addition to the matrix file. Panache can be used to display graph based pangenomes if they are previously linearized with tools such as *BioGraph.jl* (https://github.com/nguyetdang/BioGraph.jl). Example files for trial and details about how to format datasets are provided on the GitHub repository.

## Discussion

Panache offers an innovative web interface for the linear representation of pangenomes, explorable through a web interface, making the exploration and use of pangenomes easier. As a lightweight application, it can be embedded easily as an iframe to complement other interfaces in existing genome information systems. It has already been tested with open access data sets (Golicz, Bayer, *et al.*, 2016; Rijzaani *et al.*, 2021) and proved to be an effective alternative to existing tools by highlighting inherent pangenomic properties. However, as a visualization tool, results are highly dependent on the methods used to create and linearize the pangenomes. Further work on pangenome representations, and particularly on structural variations, is still needed to enhance existing representations.

Current plans for Panache include native support of graph files such as *GFA* (https://github.com/GFA-spec/GFA-spec) as an input and faster display technologies like WebGL. Panache is still under active development and new features of interest to the community will be added regularly through its GitHub page.

## Acknowledgements

The authors wish to thank Romain Basset and Mel Florance for their help on Vue.js and improvements brought to the code, as well as Dr. Eric Ganko and Steve Graham for their useful feedback. The authors acknowledge the ISO 9001 certified IRD itrop HPC (member of the South Green Platform) at IRD montpellier for providing HPC resources that have contributed to the research results reported within this paper. URL: https://bioinfo.ird.fr/-http://www.southgreen.fr.

## Funding

This work was supported by funding from Agropolis Fondation’s ‘GenomeHarvest’ project (ID 1504-006), CIFRE doctoral contract 2018/1475, Syngenta and the CGIAR Research Program, Roots, Tubers and Bananas.

The authors declare to have no conflict of interest.

## Notes

### Competing Interest Statement

The authors have declared no competing interest.

### Summary of Updates

Error in the author names, Francois Sabot has no middle name.

## References

Beyer,W. et al. (2019) Sequence tube maps: making graph genomes intuitive to commuters. Bioinformatics, 35, 5318–5320.

Computational Pan-Genomics Consortium, (2018) Computational pan-genomics: status, promises and challenges. Brief Bioinform, 19, 118–135.

Ding,W. et al. (2018) panX: pan-genome analysis and exploration. Nucleic Acids Research, 46, e5–e5.

Garrison,E. et al. (2018) Variation graph toolkit improves read mapping by representing genetic variation in the reference. Nature Biotechnology, 36, 875–879.

Golicz,A.A. et al. (2019) Pangenomics Comes of Age: From Bacteria to Plant and Animal Applications. Trends in Genetics.

Golicz,A.A., Bayer,P.E., et al. (2016) The pangenome of an agronomically important crop plant Brassica oleracea. Nat Commun, 7, 13390.

Golicz,A.A., Batley,J., et al. (2016) Towards plant pangenomics. Plant Biotechnol. J., 14, 1099–1105.

Hennig,A. et al. (2015) Pan-Tetris: an interactive visualisation for Pan-genomes. BMC Bioinformatics, 16, S3.

Li,H. et al. (2020) The design and construction of reference pangenome graphs with minigraph. Genome Biology, 21, 265.

Maston,G.A. et al. (2006) Transcriptional Regulatory Elements in the Human Genome. Annual Review of Genomics and Human Genetics, 7, 29–59.

Nielsen,C. and Wong,B. (2012) Representing the genome. Nature Methods, 9, 423–423.

Pedersen,T.L. et al. (2017) PanViz: interactive visualization of the structure of functionally annotated pangenomes. Bioinformatics, 33, 1081–1082.

Rijzaani,H. et al. (2021) The pangenome of banana highlights differences between genera and genomes. The Plant Genome, (in press).

Sherman,R.M. et al. (2019) Assembly of a pan-genome from deep sequencing of 910 humans of African descent. Nature Genetics, 51, 30–35.

Tranchant-Dubreuil,C. et al. (2019) Plant Pangenome: Impacts on Phenotypes and Evolution. In, Annual Plant Reviews online., pp. 453–478.

Wick,R.R. et al. (2015) Bandage: interactive visualization of de novo genome assemblies. Bioinformatics, 31, 3350–3352.

Yoghourdjian,V. et al. (2020) Scalability of Network Visualisation from a Cognitive Load Perspective. IEEE transactions on visualization and computer graphics, PP.

Yokoyama,T.T. et al. (2019) MoMI-G: modular multi-scale integrated genome graph browser. BMC Bioinformatics, 20, 548.

